# Meta-Analysis of Rumen-Native Microbial Supplementation on Culling Risk and Herd Retention in Commercial Dairy Herds

**DOI:** 10.64898/2026.10.01.755474

**Authors:** Clarisse Marotz, Thys Jacobi, Josh Lefler, Brooke Anderson, Ivan Zhelev, Savannah Katulski, John Goeser, Mallory Embree

**Affiliations:** Native Microbials, San Diego, CA 92101; Department of Animal and Dairy Sciences, University of Wisconsin-Madison, WI 53706

**Keywords:** rumen microbiome, rumen-native microbes, direct-fed microbial, dairy cattle, culling risk, herd retention, longevity, sustainable dairy production

## Abstract

**Background:** Premature culling reduces dairy farm profitability and increases the environmental footprint of milk production by shortening productive lifespan and increasing replacement animal requirements. This study evaluated whether treatment with a live consortium of rumen-native microbes was associated with improved herd retention across controlled commercial dairy trials. Cull outcomes (sold or died vs. retained in herd) were analyzed across twenty trials encompassing more than 40,000 dairy cattle housed across 110 experimental pens, with each trial conducted for at least 5 months. Treatment pens received a defined daily dose of Galaxis® Frontier (“GF”, Native Microbials, San Diego, CA), containing *Pichia kudriavzevii*, *Clostridium beijerinckii*, *Ruminococcus bovis*, and *Butyrivibrio fibrisolvens,* incorporated into the ration; control and treatment groups were otherwise managed equivalently. Cow-level time to culling was modeled within each trial, accounting for pen-level clustering, and trial estimates were combined using random-effects meta-analysis.

**Results:** Supplementation with rumen-native microbes was associated with a 9% lower hazard of culling across 20 trials (hazard ratio = 0.91, 95% confidence interval: 0.86-0.96; P < 0.001). Between-trial heterogeneity was negligible, and treatment effects were not significantly moderated by breed or geographic region. Detailed health records from one trial were examined to further characterize health outcomes associated with GF; treatment was associated with fewer transition-related disorders, particularly ketosis and metritis, whereas mastitis and lameness were largely unaffected. These findings suggest that the health effects associated with GF may be more pronounced for disorders linked to physiological and metabolic adaptation during the transition period than for health events more broadly.

**Conclusion:** Across diverse commercial dairy environments, feeding a defined consortium of rumen-native microbes was associated with improved dairy cow retention. Recorded health events suggested that improved metabolic stability may contribute to this effect. Extending productive lifespan reduces replacement requirements and the number of non-lactating heifers needed to maintain herd size, thereby reducing the economic burden and carbon footprint associated with replacement animals that do not directly contribute to milk production. These results demonstrate rumen-native microbial supplementation as a practical strategy to enhance the economic and environmental sustainability of dairy production.

## Background

Improving productive lifespan in dairy cattle is critical not only for animal welfare, but also for dairy farm economic performance and environmental sustainability. Cows that remain in the herd longer can produce more lifetime milk, reduce replacement requirements and costs, and decrease the environmental footprint of milk production by reducing the number of non-lactating replacement animals required to maintain herd size [1]. While many factors influence herd longevity, a cow’s ability to respond to and endure metabolic, inflammatory, or infectious challenges plays a central role [2]. Acute immune system activation alone can consume >1 kg of glucose in 12 hours [3], the dietary equivalent of several kilograms of corn grain. Cows with better health and energy status are more likely to recover from the transition period, successfully rebreed, sustain milk production and remain in the herd. Therefore, identifying strategies that enhance baseline metabolic and immune capacity is essential to building longer-lasting herds and improving feed conversion efficiency.

In ruminants, metabolic capacity is dictated largely by the diverse microbial ecosystem native to the gastrointestinal tract, with estimates that up to 70% of a cow’s energy demands are supplied entirely by the rumen microbiome [4]. Because rumen microorganisms function as an interconnected community, changes in key microbial populations can influence the functional capacity of the broader ecosystem. Galaxis® Frontier (GF) contains four rumen-native organisms, *Pichia kudriavzevii*, *Clostridium beijerinckii*, *Ruminococcus bovis*, and *Butyrivibrio fibrisolvens*, which were identified through broad sequencing surveys of productive dairy cows, followed by network analyses that linked specific microbial population with animal metadata and productive performance [5].

Recent work has highlighted the reproducible ability of GF to improve feed efficiency [6–13] and increase milk production in animals when GF was fed starting in early lactation [6–11]. These responses are accompanied by changes in rumen fermentation, including shifts in the ruminal volatile fatty acid profile [8–9], as well as downstream changes in host physiology. Tabor et al. reported that cows supplemented with GF exhibited increased neutrophil activity and greater mammary tissue capacity [13] and Bulnes et al. identified an increased acute phase response during the transition to lactation that was later followed by increased milk production [8]. Together, these findings indicate that the effects of rumen-native microbial supplementation extend beyond ruminal fermentation and are associated with systemic immunometabolic responses.

To evaluate whether these systemic effects of rumen-native microbial supplementation translate into meaningful health and performance outcomes, we focused on culling, a key determinant of both herd profitability and sustainability. Premature culling is most often driven by health and performance challenges, including low milk production, metabolic and infectious diseases, reproductive failure, and lameness [14].

Thus, cull rate serves as an integrative endpoint, reflecting the cumulative impact of resilience on cow survival and productive longevity. Importantly, culling status (sold or died) represents a binary, unambiguous outcome that is consistently recorded across commercial farms enabling robust comparison across diverse production systems. Because health event definitions and recording practices differ among herds, cull data offer a more standardized and auditable outcome for large-scale field analyses. Here, we report results from a multi-site treatment-versus-control analysis conducted in U.S. commercial dairy herds.

In late 2023, high-pathogenicity avian influenza (HPAI) spilled over into U.S. dairy cattle and spread across multiple states [15]. HPAI outbreaks altered herd management and, in some herds, increased culls independent of underlying cow health [16]. Because several study herds experienced documented HPAI outbreaks during their trial periods, HPAI status was evaluated as a potential moderator of treatment effect and affected trials were identified in the primary meta-analysis. Finally, to further characterize health outcomes associated with GF supplementation, we examined detailed health-event records from one commercial trial.

## Methods

### Trial Design and Data Collection

#### Trial farm enrollment

Commercial dairies meeting the following criteria were selected as trial sites: 1) ear tags that allow tracking daily pen location for each cow, 2) dairy cow management software tracking individual cow location, fresh dates, dry dates, and cull date, 3) two or more pens with matched management and demographics; pens were considered ‘matched’ when animals had similar averages and distributions of parity (lactation number), days in milk, and baseline milk production, thereby ensuring comparability between treatment and control groups, and 4) capability to deliver treatment consistently to designated pens.

#### Trial design

At each trial site, matched pens were selected based on parity, days in milk, and baseline milk production. Matched pens were then randomly assigned to either the control group (base total mixed ration, or TMR, using the current farm formulation) or the treatment group (“GF”; fed base TMR with the addition of Galaxis® Frontier). The treatment pens received 5 g/head/day GF, which contains *Clostridium beijerinckii* ASCUSDY20 at 1.0 × 10^7^ colony forming units (CFU), *Pichia kudriavzevii* ASCUSDY21 at 1.0 × 10^8^ CFU, *Ruminococcus bovis* ASCUSDY10 at 1.0 × 10^8^ CFU, and *Butyrivibrio fibrisolvens* ASCUSDY19 at 1.0 × 10^8^ CFU. Pens remained open during the trial, allowing continuous enrollment as new cows entered the pen. A power analysis, based on pen enrollment rates and weekly cow-level milk variance, was used to determine the length of trial required to enroll enough cows to detect a 1.36 kg (3 lbs) cow-level milk yield improvement when comparing the lactation curves of cows that enrolled pre-peak.

#### Microtracer auditing

To confirm adequate mixing and delivery of the full dose of GF (5 grams/head/day), a Microtracers™ F Blue-lake #1 tracer (Micro-Tracers, San Francisco, CA) was used to audit total mixed ration (TMR) preparation at each trial site prior to the trial initiation according to the manufacturer’s instructions. Briefly, 50 grams of inert iron particles with colorant was added per ton of TMR, directly to the mixer wagon and at the same time as GF. Ten samples were collected along the bunk in order of feed-out and sent to Native Microbials for magnetic separation and quantification. Sites that had >80% recovery of iron particles and a coefficient of variation <20% across the ten samples were approved to proceed with the trial.

#### Data collection

Automatic herd management software data exports were set up at each trial site, resulting in daily CSV files with each cow’s ID, days in milk, lactation number, pen location, and production data (either test day milk yields or daily milk meter weights as available). At the end of the trial, a report was generated from the herd dairy software providing fresh dates, dry dates, and cull (sold/died) dates when applicable for each animal enrolled in the trial.

#### Data Processing

A standardized quality control (QC) process was applied to each dataset to ensure each cow was assigned to a single treatment group throughout the trial. Daily tracking data from each site was compiled into a single CSV dataframe. For each cow, the first appearance in a trial pen on or after the trial start date (either treatment or control) was labeled as day_cow=1, with subsequent days numbered sequentially. If a cow experienced multiple lactations during the trial period (i.e. dried off, calved and re-entered a trial pen), only the lactation with the longest trial enrollment was retained for analysis. To prevent any unintentional animal movement from confounding results, cows were unenrolled after being found in the wrong treatment group; specifically, a cow that switched between treatment and control pens for more than five total days had all data after their fifth switched day removed. If a cow was missing from the daily export data for more than five consecutive days, her data was excluded from that point onward.

Once the QC process was complete, the retained data was merged with calving and culling records to determine the number of days each cow remained in the trial and whether she was culled, providing the inputs for the survival analysis.

For the survival analysis, each cow was represented by data on her final observed day in the trial. For cows that were culled, this corresponds to the date of culling. For cows that were not culled, the data reflects the last day she was observed in the trial during the trial period. To focus on animals that are the most common target of culling decisions and improve the consistency of the dataset, the following inclusion criteria were applied. First, cows with >400 days in milk at trial enrollment were excluded, as these animals were more likely to be scheduled for culling regardless of product effect. Second, only cows that remained in the trial for at least 8 weeks (56 days) were included, ensuring sufficient exposure to the product. Finally, to account for farm logistics such as holding pens, cows were allowed to be classified as culled up to 30 days after exiting a trial pen.

#### Trial Inclusion

We initiated analyses with a total of 37 treatment vs control trials. Four were excluded a priori for protocol or measurement errors (incorrect dosing, missing sold/died data, sold/died records corrupted by a herd-file merge, or prematurely ended due to systematic parlor-based milk-test bias). Trials with insufficient sold/died events to detect a ≥50% reduction in culls with 70% power (α=0.08) were removed (n=4). Baseline differences between treatment groups were then evaluated for milk yield, lactation number, and days in milk (DIM), as each is an established predictor of culling risk [14]. Milk yield was averaged over each cow’s first 7 days following enrollment. To minimize confounding, trials were excluded if treatment and control groups were mismatched in at least two of the three covariates, defined as both Mann-Whitney U P < 0.05 and |Cohen’s d| > 0.2 for each covariate (n=5), or if any single covariate showed substantial imbalance (|Cohen’s d| > 0.5), regardless of P-value (n=3). These screening criteria resulted in 21 eligible trials. Because treatment was assigned at the pen level, trials included in the primary analysis were additionally required to have replication of the experimental unit within each treatment arm (i.e. ≥2 pens assigned to each of treatment and control). One trial had only 1 treatment and 1 control pen and was therefore excluded because treatment effects could not be distinguished from pen-level effects. The final dataset consisted of 20 trials comprising 42,107 animals; a more detailed breakdown of trial-level covariate metrics can be found in Supplemental Table 1.

### Statistical Analyses

All analyses were performed using R (R Core Team, 2020) (v4.4.1) and tidyverse package suite (v2.0.0)[17].

#### Mixed-effects Cox proportional hazards models

Time to culling was analyzed separately within each trial using mixed-effects Cox proportional hazards models implemented with the *coxme* package (v2.2.22) [18]. Treatment was applied at the pen level, therefore pen was considered the experimental unit and individual cows the observational units. Because cows move among pens on commercial dairies, each cow was assigned a primary experimental pen for the survival analysis. For each cow, the number of days spent in each trial pen was counted, and the pen occupied for the greatest number of days was designated as the cow’s primary pen. The model included treatment group, lactation group, and days in milk at trial enrollment as fixed effects, with a random intercept for the cow’s primary experimental pen. Lactation was modeled categorically (1, 2, 3+), and DIM at enrollment was modeled as a continuous covariate. The model for each trial was specified as:

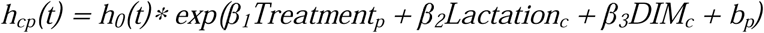

where *h_cp_(t)* is the culling hazard for cow *c* in pen *p* at day *t*, *h_0_(t)* is the baseline hazard, β*_1_* is the treatment effect, and *b_p_* is the random effect for pen. Pen random effects were assumed to follow a mean-zero Gaussian distribution. Thus, inference for the treatment effect accounted for the shared pen-level clustering arising from assignment of treatment at the pen level.

The treatment effect from each trial was expressed as a hazard ratio (HR), calculated as exp(β*_1_)*, where β*_1_* is the estimated Cox coefficient for treatment on the log-hazard scale. The HR represents the relative instantaneous hazard of culling for GF-treated cows compared with control cows, conditional on the fixed effects and pen-level random effect. An HR < 1 indicates a lower culling hazard in the GF group. For interpretability, the relative reduction in culling hazard was calculated as (1 – HR) * 100%; for example, an HR of 0.87 corresponds to a 13% lower estimated culling hazard in GF-treated cows relative to controls.

#### Meta-analysis of trial-level effects

To synthesize findings across farms, trial-specific HR estimates from the mixed-effects Cox models were pooled using a random-effects meta-analysis via the *meta* package (v7.0-0) [17]. Trial weights were assigned by inverse variance, incorporating both the within-trial variance of the treatment estimate and between-trial variance (τ²). Between-trial variance was estimated using the Paule-Mandel estimator, with confidence intervals for the pooled effect calculated using the Hartung-Knapp method. Between-trial heterogeneity was assessed using Cochran’s Q statistic and I² [19–21]. Trial-specific and pooled effects were reported as HR with 95% confidence intervals.

#### Moderator analyses

To evaluate whether treatment effects varied across trial characteristics, separate meta-regression models were fit for breed group (Holstein vs. non-Holstein), geographic region (Western vs. Midwestern), and HPAI status (affected vs. unaffected during the trial period) using meta::metareg. Meta-regression was performed on the trial-level treatment effects from the primary random-effects meta-analysis. Subgroup-specific hazard ratios were estimated from the fitted meta-regression coefficients and transformed from the log-hazard-ratio scale to the hazard-ratio scale. Moderator *P*-values tested whether treatment effects differed between moderator categories.

#### Event-specific hazard analysis

An exploratory analysis of cow-level health events was performed for one commercial herd (CAH01) that maintained comprehensive health-event records. The trial began on a fixed calendar date and included cows across stages of the lactation; cows already present at trial initiation entered the study at their current stage of lactation, whereas subsequent enrollment occurred as cows entered designated treatment or control close-up pens starting 3-4 weeks before their predicted calving date. Treatment and control groups were continued in designated pens through lactation, allowing health outcomes to be evaluated across the transition period and later lactation. Health outcomes were evaluated individually, including mastitis, metritis, ketosis, milk fever, lameness, diarrhea, blood in milk, infection, and as pooled categories. Pooled transition-associated events included metritis, ketosis, or milk fever and pooled non-transition associated events included mastitis, lameness, diarrhea, blood in milk, or infection.

Because transition events are tightly linked to physiological stage, time to event was measured in days in milk (DIM) rather than days enrolled in the trial (day_cow). For transition events, follow-up was limited to the first 60 DIM, when these conditions are most likely to occur. For non-transition events, cows were followed through the remainder of lactation. Observation started at trial enrollment and ended at the earliest of: (i) the first occurrence of the event of interest, (ii) culling or sale, or (iii) the trial end date. Only the first occurrence of the event(s) of interest was analyzed. Cows that did not experience the event were considered event free up to their last recorded observation (or at 60 DIM for transition events). Events recorded before trial start were excluded.

Adjusted survival curves were generated from the fitted Cox proportional hazards models and displayed as cumulative event probability (1 − *S(t)*) with 95% confidence intervals. Treatment effects were evaluated using cow-level Cox proportional hazards models including treatment (GF vs. Control) and parity category (lactation number: 1, 2, 3, 4, ≥5) as fixed effects. Hazard ratios (HR) and 95% confidence intervals are reported for GF relative to Control, with relative hazard reduction calculated as (1 − HR) × 100%. Treatment *P*-values were obtained using likelihood-ratio tests comparing the full model with a reduced model excluding treatment. All tests were two-sided with α = 0.05.

Treatment was assigned at the pen level during both the close-up and lactating periods. However, individual cow assignments to replicated close-up pens were not retained in the herd-management software, preventing retrospective reconstruction of the pen-level clustering for transition events. Because most transition events occurred during or shortly after this period, event-specific health analyses were therefore conducted at the cow level and should be considered exploratory.

## Results

### Trial characteristics

Across the 20 commercial dairy trials included in the meta-analysis (Table 1), study populations spanned eight U.S. states (CA, WI, WA, SD, ID, OH, MN, and MI) and included predominantly Holstein herds (n=14), with additional Jersey (n=3) and crossbred (n=3) populations. Trial duration ranged from 21 to 43 weeks (median, 28 weeks; ∼7 months) and encompassed 42,107 animals (20,715 Control and 21,392 GF).

**Table 1.** Baseline covariates by treatment across trials. Values are sample-size–weighted means for each arm (Control, GF); “±” denotes the between-trial SD of arm-level means, weighted by arm size. Covariates were calculated from each cow’s first week after enrollment and averaged per trial. Parenthetical ranges are unweighted trial-level min-max.

|  | Control | GF |
| --- | --- | --- |
| **Days in Milk** | 109 ± 25 [63–157] | 104 ± 24 [60–160] |
| **Lactation** | 2.86 ± 0.61 [2.04–4.03] | 2.81 ± 0.54 [1.87–4.14] |
| **Baseline Milk Yield (lbs)** | 91.04 ± 12.60 [67.92–110.91] | 92.56 ± 13.53 [68.59–113.69] |
| **% 1st lactation** | 17.6 ± 16.7 [0.0–45.6] | 15.7 ± 16.2 [0.0–49.6] |
| **Number Pens** | 3 ± 1 [2–7] | 3 ± 1 [2–6] |
| **Total cows (N)** | 20,715 | 21,392 |

Because trials began on fixed calendar dates, enrolled populations included both cows already present in study pens at trial initiation and cows enrolled through close-up pens before calving. Average DIM at enrollment varied widely among sites, from ∼60 DIM in WIH02, where enrollment began during the fresh period, to >150 DIM in MNH01 and MIH02, which included later-lactation pens. Most trial populations averaged 90–130 DIM at enrollment. Average parity was between 2 and 3 in most trial populations, while the proportion of first-lactation cows ranged from near 0% to >45%. Baseline milk production averaged approximately 95 ± 10 lb/d among Holstein groups and 77 ± 10 lb/d among Jersey and crossbred groups. Treatment groups included approximately 400–2,400 observational units per arm, distributed across 2–7 experimental pens per treatment arm.

### Overall Meta-analysis results

Control group cull rates during the trial period ranged from 5.20% to 13.48% (average 8.46%) across the 20 trial sites. Random-effects meta-analysis of treatment effects produced a pooled hazard ratio (HR) of 0.91 (95% CI 0.86 – 0.96; p = 0.0008), corresponding to a 9% lower hazard of culling in the treatment group versus control (Figure 1). Between-trial heterogeneity was negligible (Cochran’s Q = 4.82, df = 19, p = 0.99; I² = 0% [0–48%]; τ² = 0).

**Figure 1.**
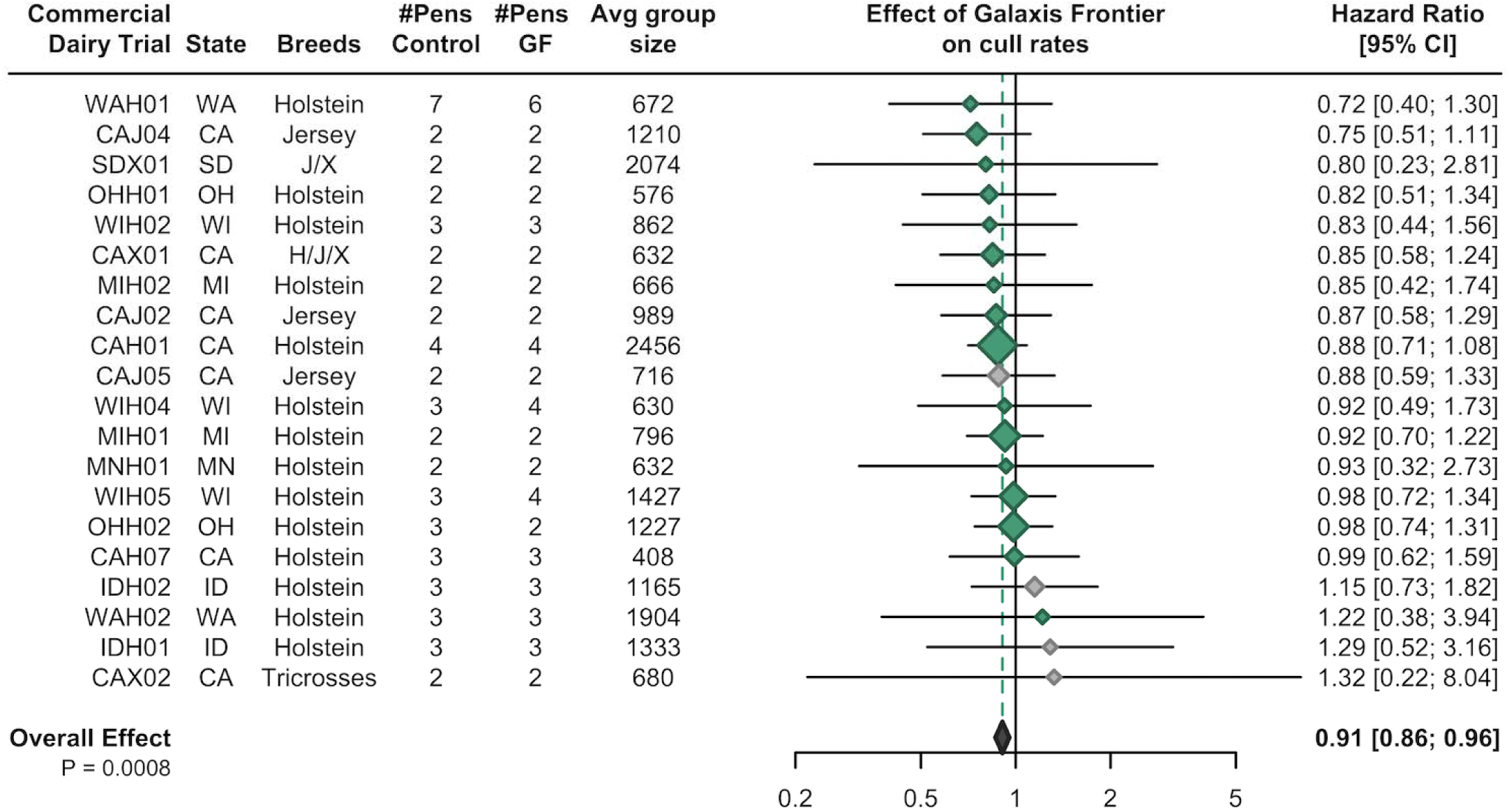
Hazard ratios for culling across 20 commercial dairy trials. Hazard ratios (HR) less than 1 indicate a low r culling hazard in the GF-treated group relative to control. Relative hazard reduction was calculated as *(1 − HR) × 100%*. Marker size reflects the relative precision of each trial estimate, with larger diamonds representing more precise estimates and greater contribution to the pooled meta-analysis. Trials shown in gray experienced HPAI outbreaks during the trial period; trials shown in green did not.

A leave-one-out sensitivity analysis confirmed the result was not driven by any one trial; the pooled relative hazard reduction ((1-HR)*100) from any 19 of the 20 studies ranged from 8.2% to 11.9% (p ≤ 0.005).

### Moderator analyses; examining the impact of breed, region, and disease status

Meta-regression was used to evaluate whether treatment effects varied according to breed composition, geographic region, or occurrence of HPAI during the trial period. Effect estimates differed numerically between breed groups, although the breed moderator did not reach statistical significance (P=0.065) with a larger estimated reduction in culling hazard among non-Holstein cattle (Holstein HR=0.93, 95% CI 0.88-0.98; non-Holstein HR=0.84, 95% CI 0.76-0.92). There was no evidence of a modifying effect by region (P=0.337; Western trial HR=0.89, 95% CI 0.83-0.95; Midwestern trial HR=0.93, 95% CI 0.87-1.0). HPAI-unaffected trials had an estimated HR of 0.89 (95% CI 0.85-0.94) compared with 1.03 (95% CI 0.90-1.17) among HPAI-affected trials, with HPAI status showing a tendency to modify the treatment effect (*P* = 0.076). Three of the four trials with HR > 1 occurred at sites experiencing HPAI outbreaks during the trial period.

### Event-specific hazard analysis at a high-fidelity site (CAH01)

To explore health outcomes that may contribute to the observed difference in herd retention, we examined detailed cow-level health records from CAH01 (Figure 2). GF treatment was associated with a 28% lower hazard of transition-related health events (HR = 0.72, 95% CI: 0.57–0.90; *P* = 0.004). Among individual transition disorders, ketosis hazard was 48% lower (HR = 0.52, 95% CI: 0.29–0.94; *P* = 0.025) and metritis hazard was 26% lower (HR = 0.74, 95% CI: 0.57–0.97; *P* = 0.029) in the GF group after adjustment for parity. Milk fever did not differ significantly between treatment groups.

**Figure 2.**
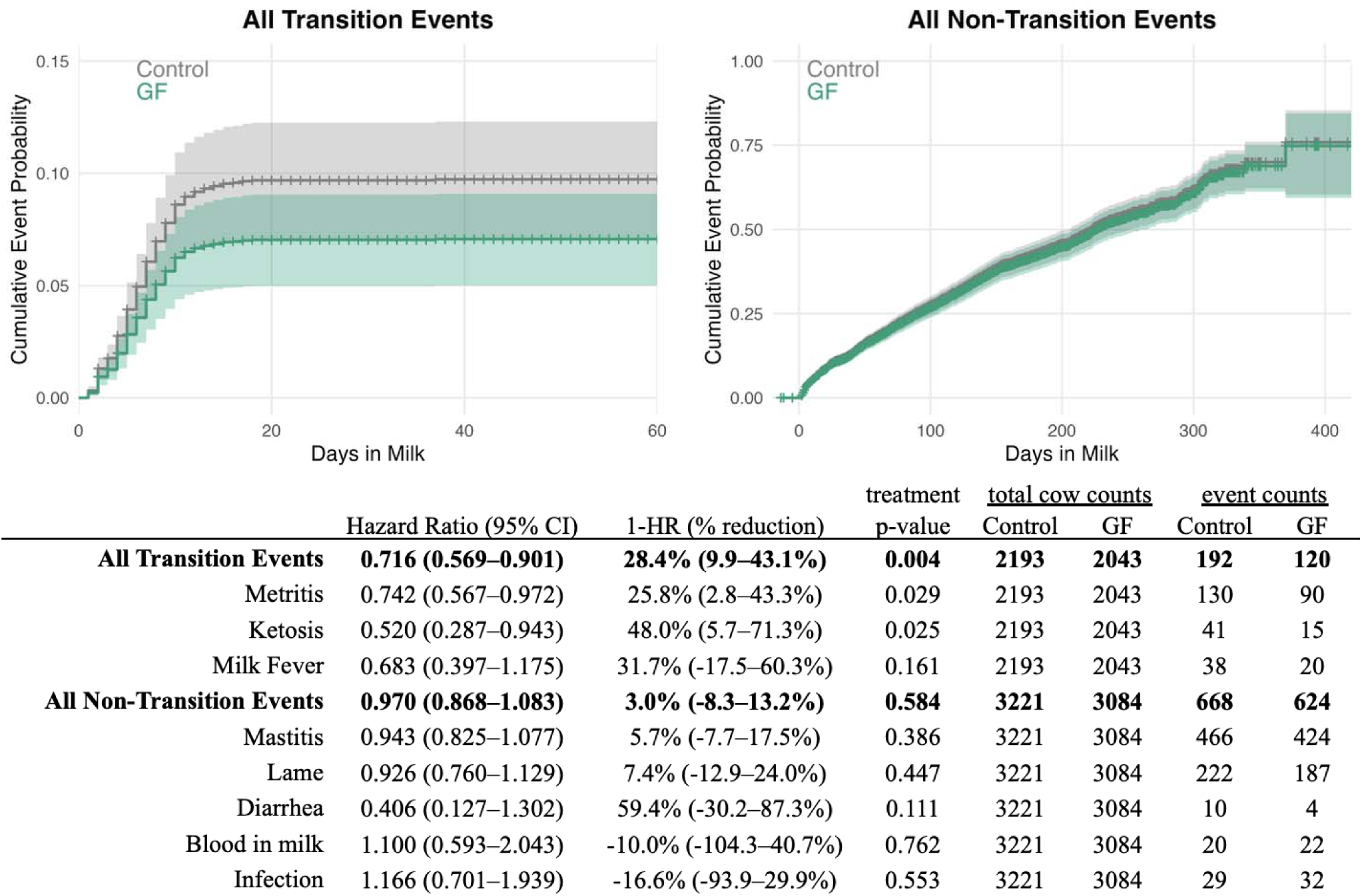
Adjusted cumulative event probability for common health events tracked at CAH01. Cox proportional hazards models were adjusted for parity (lactation group). Curves display the model-estimated cumulative event probability, *1 − S(t)*, with 95% confidence intervals. Hazard ratios (HR) compare the hazard of an event in GF-treated cows with that in control cows; HR < 1 indicates a lower event hazard in the GF group. Relative hazard reduction was calculated as *(1 − HR) × 100%*, with confidence bounds derived from the HR 95% confidence interval. Treatment *P*-values were obtained from likelihood-ratio tests comparing the full model with a reduced model excluding treatment. Transition-event analyses were restricted to ≤60 days in milk.

In contrast, associations with non-transition health events were modest. Mastitis and lameness, with 14% and 7% incidence respectively in the control group, were the most common non-transition events and were not significantly different in the GF group. Diarrhea occurred infrequently (<2%) and blood in mil and infection were rare; none of these health events showed evidence of a treatment effect. Overall, the hazard of pooled non-transition events did not differ between experimental groups (HR = 0.97, 95% CI: 0.87–1.08; *P* = 0.584).

## Discussion

To our knowledge, this is the first study to evaluate the impact of a rumen-native microbial product or direct-fed microbial on cull rates in controlled commercial trials. Across 20 trials encompassing 42,107 cows, daily GF supplementation was associated with a 9% lower hazard of culling, and the pooled effect remained significant in a sensitivity analysis of the impact of individual trials. Despite substantial diversity in geography, breed, parity structure, management, and stage of lactation at enrollment, there was little evidence of between-trial heterogeneity beyond that expected from sampling variation. Together, these findings suggest that targeted manipulation of the rumen microbiome can translate beyond changes in rumen fermentation and production efficiency to measurable differences in herd retention under commercial conditions. These effects are consistent with the broader biological role of the rumen microbiome, which not only drives feed digestion and fermentation end products, but also modulates immune signaling and nutrient partitioning [22], providing plausible pathways through which changes in the rumen microbial community could influence longer term cow health and retention.

Prior studies supplementing rumen-native microbes have demonstrated effects on ruminal fermentation, nutrient utilization, and host metabolic and immune measures [7–9, 13, 23]. By contrast, conventional microbial feed supplements composed of non-native organisms (e.g., *Saccharomyces cerevisiae*, *Bacillus subtilis)* have produced heterogeneous effects on systemic immunometabolic biomarkers during the transition period [24–26]. This divergence raises the possibility that ecological compatibility of rumen-native microorganisms with the resident rumen ecosystem may contribute to the consistency of the observed responses across farms. Most prior studies testing the effects of GF have been conducted in small academic cohorts, typically involving only tens of cows per group. These smaller studies provide important physiological insight but do not readily translate to herd-level outcomes. In contrast, commercial-scale trials allow health effects to be evaluated under real-world conditions, where impacts on survival and culling risk can be directly measured. Control-group culling averaged 8.46% during the study periods, lower than annual herd-level culling rates commonly reported for U.S. dairy herds, which often range from 30-40% [1]. This difference can be explained in part because the present analysis evaluated only trial-eligible animals over substantially shorter observation periods rather than complete annual herd turnover. Nevertheless, the large population and diversity of participating herds allowed the treatment effect to be evaluated across a broad range of commercial production environments.

The magnitude of the treatment effect was not significantly moderated by geographic region, breed, or herd HPAI status. However, moderator analysis provided suggestive evidence of variation by both breed (*P* = 0.065) and HPAI status (p=0.076). One potential explanation for the larger estimated effect in non-Holstein herds is relative dose, as GF was administered at a fixed dose per cow rather than adjusted for body weight; this hypothesis warrants evaluation in future dose-response studies.

HPAI represented an important additional source of real-world disruption during the study period. Three of the four trials with HR > 1 experienced documented HPAI outbreaks, and the estimated treatment effect was attenuated among HPAI-affected trials (HR = 1.03) relative to unaffected trials (HR = 0.89). Acute infectious disease outbreaks can alter both cow health and herd-management decisions, potentially obscuring treatment-associated differences in routine culling. Because these changes affected all cows, they may have diluted the relative treatment effect and introduced new sources of variation during the trial. We chose to retain these trials in the primary analysis and report the observable effects of GF across the full range of commercial conditions encountered during the study.

Cull-event logging practices varied across farms, particularly with respect to recording reasons for culling, as is typical in large commercial settings where multiple individuals may record events using different definitions. By contrast, whether a cow was removed from the herd (sold or died) is a binary outcome that is consistently and reliably captured across all sites, providing a robust basis for this analysis. To explore health outcomes that might contribute to herd retention, detailed event records from CAH01 were analyzed separately. GF supplementation was associated with lower hazards of transition-related disorders, particularly ketosis and metritis, whereas there were no associations identified between GF supplementation and non-transition-associated events. Because transition disorders are closely linked to energy balance and metabolic adaptation around parturition [27], this pattern is consistent with the hypothesis that improved ruminal metabolism and metabolic stability during the transition period may contribute to improved herd retention.

Several limitations should be considered when interpreting these findings. Although treatment was applied at the pen level and trial-specific models accounted for clustering using each cow’s predominant experimental pen, cows could move among pens during commercial management and complete longitudinal pen histories could not be incorporated into the model. Commercial trials also necessarily differed in duration, enrollment stage, management practices, and underlying populations. While all-cause removal provides a robust endpoint across farms, it combines voluntary and involuntary culling decisions and therefore cannot distinguish whether improved herd retention was more likely due to management decisions versus health-based reasons. Finally, the CAH01 health-event analysis was exploratory. Individual assignments to replicated close-up pens were not retained in herd-management records, preventing adjustment for close-up pen-level clustering and limiting causal inference from these event-specific outcomes. Future prospective studies with more stringent pen or treatment management and standardized health-event definitions will be valuable for confirming which physiological and health pathways mediate the observed improvement in herd retention.

Taken together, these findings demonstrate a reproducible association between feeding a defined rumen-native microbial consortium and improved herd retention across diverse commercial dairy environments. The concentration of health-associated differences around the transition period provides a plausible link between rumen microbial function, transition period health, metabolic resilience, and productive longevity that warrants prospective investigation. By reducing premature removal and extending productive lifespan, microbiome-targeted strategies may reduce replacement requirements and improve the economic and environmental efficiency of dairy production.

## Conclusions

Across 20 controlled commercial dairy trials encompassing 42,107 cows housed across 110 pens, feeding a defined consortium of rumen-native microbes was associated with a reproducible reduction in culling hazard across diverse production environments. Complementary health-event data suggest that improved transition period health and metabolic stability may contribute to this effect. These findings demonstrate that targeted modulation of the rumen microbiome can translate into meaningful herd-level outcomes under commercial conditions. By improving herd retention and productive longevity, rumen-native microbial treatment represents a practical microbiome-targeted strategy to reduce replacement requirements and improve the economic and environmental sustainability of dairy production.

## Supporting information

Supplemental Table 1; trial metadata

## List of abbreviations

DIM: Days in milk; day of calving was defined as DIM 0
GF: Galaxis® Frontier
CFU: Colony forming unit
HPAI: Highly pathogenic avian influenza
HR: Hazard ratio
VFA: Volatile fatty acid
TMR: Total mixed ration

## Declarations

### Ethics approval and consent to participate

The data analyzed in this study originated from commercial dairy operations under routine management conditions, including feeding of a registered feed additive. No procedures outside normal commercial husbandry and management practices were performed for the purposes of this study. Data were collected with producer consent and in accordance with applicable animal welfare requirements.

### Consent for publication

Not applicable

### Availability of data and materials

All analysis scripts and outputs are publicly available on Github [https://github.com/lisa55asil/Galaxis-Frontier-survivability-meta-analysis]. The underlying anonymized row-level farm data are not publicly posted due to privacy considerations but can be made available upon reasonable request to the corresponding author.

### Author contributions

Clarisse Marotz, Brooke Anderson, Josh Lefler, and Mallory Embree contributed to study design. Savannah Katulski and Ivan Zhelev oversaw and managed study implementation and data collection. Statistical analyses were performed by Thys Jacobi, Clarisse Marotz, Josh Lefler, and Brooke Anderson. Clarisse Marotz drafted the manuscript. John Goeser, Mallory Embree, Brooke Anderson, Josh Lefler and Savannah Katulski contributed to manuscript editing and revision. All authors reviewed and approved the final manuscript.

## Acknowledgement

Appreciation is extended to Dr. Jon Robison for guidance in experimental design and data interpretation.

---

**Supplemental table 1.**
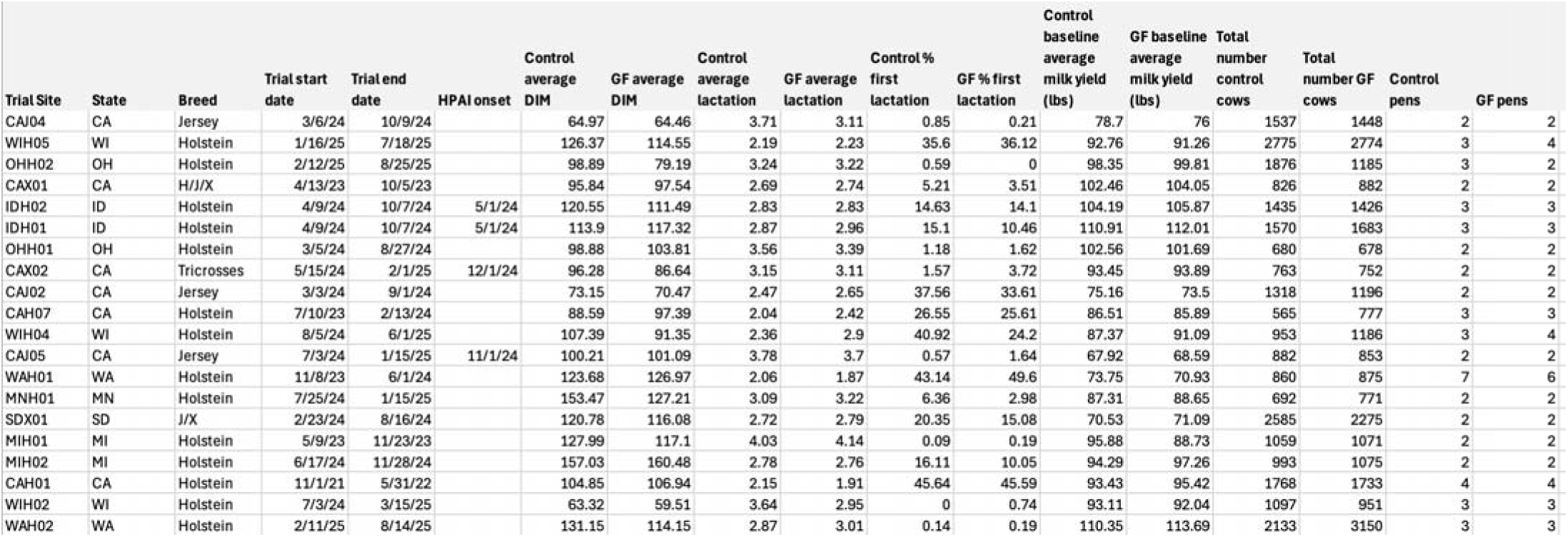
Trial-level covariate metrics. Trial details including location (state), breeds enrolled, trial start and end dates, HPAI outbreak onset (where applicable), and mean days in milk (DIM), lactation number, and milk yield during each cow’s first week of trial enrollment.

## References

1. De Vries A. Symposium review: Why revisit dairy cattle productive lifespan?.Journal of Dairy Science. 2020 Apr 1;103(4):3838–45.

2. Sordillo LM, Contreras GA, Aitken SL. Metabolic factors affecting the inflammatory response of periparturient dairy cows. Animal health research reviews. 2009 Jun;10(1):53–63.

3. Kvidera SK, Horst EA, Abuajamieh M, Mayorga EJ, Fernandez MS, Baumgard LH. Glucose requirements of an activated immune system in lactating Holstein cows. Journal of dairy science. 2017 Mar 1;100(3):2360–74.

4. Mizrahi I, Jami E. A method to the madness: disentangling the individual forces that shape the rumen microbiome. EMBO reports. 2021 Feb 3;22(2):e52269.

5. Zengler K, Embree M, inventors; Ascus Biosciences Inc, assignee. Methods, apparatuses, and systems for analyzing microorganism strains from complex heterogeneous communities, predicting and identifying functional relationships and interactions thereof, and selecting and synthesizing microbial ensembles based thereon. United States patent US 9,540,676. 2017 Jan 10.

6. Valldecabres A, Gilmore SP, Embree JJ, Zhelev IZ, Gaffney JR, Marotz CA, Yang F, Izzo AS, Embree MM, Lago A. Effects of rumen-native microbial feed supplementation on milk yield, composition, and feed efficiency in lactating dairy cows. Journal of animal science. 2022 Oct 1;100(10):skac275.

7. Goldsmith K, Lefler J, Embree M, VandeHaar MJ. The effect of supplementing native rumen microbes on milk production of dairy cows. JDS communications. 2023 Jan 1;4(1):31–4.

8. Bulnes M, Bonilla J, Suazo M, Michelotti TC, Paz A, Lefler J, Marotz C, Embree M, Begalli G, Halfen J, Fernandes T. Effects of peripartal rumen-derived direct-fed microbial supplementation on lactation performance, metabolism, ruminal fermentation, and microbial abundance in dairy cows. Journal of Dairy Science. 2025 Mar 1;108(3):2354–70.

9. Marinho MN, Perdomo MC, Simões BS, Husnain A, Arshad U, Figueiredo CC, Santos JE. Dietary supplementation of rumen native microbes improves lactation performance and feed efficiency in dairy cows. Journal of Dairy Science. 2024 Oct 1;107(10):7918–31.

10. Valldecabres A, Anderson B, Lefler J, Marotz CA, Embree MM, Lago A. Impact of a native rumen microbe supplement provided in feed on production performance and enteric methane emissions in lactating dairy cows. Journal of Dairy Science. 2025 In press.

11. Ferro LN, Kerwin AL, Ryan CM, Graef GM, Nelson TM, Green H, Yang F, Embree M, Barbano DM, Overton TR. Effects of a novel direct-fed microbial feed additive on performance of lactating Holstein dairy cows. ADSA poster presentation 2022.

12. Dickerson AM, Yang F, Green HB, Embree MM, Drackley JK. Feeding native rumen microbial supplements increases energy-corrected milk production and feed efficiency by Holstein cows. JDS communications. 2022 Jul 1;3(4):239–44.

13. Tabor E, Guadagnin AR, Guenther M, Cangiano LR, Panisa F, Anderson B, Niehues L, Marotz C, Embree M, Laporta J. Effects of feeding a native rumen-native microbial live supplement during pre-and postpartum on health, performance, and blood metabolites of Holstein cows. Journal of Dairy Science. 2025 Aug 20.

14. Hadley GL, Wolf CA, Harsh SB. Dairy cattle culling patterns, explanations, and implications. Journal of dairy science. 2006 Jun 1;89(6):2286–96.

15. Nguyen TQ, Hutter CR, Markin A, Thomas M, Lantz K, Killian ML, Janzen GM, Vijendran S, Wagle S, Inderski B, Magstadt DR. Emergence and interstate spread of highly pathogenic avian influenza A (H5N1) in dairy cattle in the United States. Science. 2025 Apr 25;388(6745):eadq0900.

16. Peña-Mosca F, Frye EA, MacLachlan MJ, Rebelo AR, de Oliveira PS, Nooruzzaman M, Koscielny MP, Zurakowski M, Lieberman ZR, Leone WM, Elvinger F. The impact of highly pathogenic avian influenza H5N1 virus infection on dairy cows. Nature Communications. 2025 Jul 15;16(1):6520.

17. Wickham H, Averick M, Bryan J, Chang W, McGowan LD, François R, Grolemund G, Hayes A, Henry L, Hester J, Kuhn M. Welcome to the Tidyverse. Journal of open source software. 2019 Nov 21;4(43):1686.

18. Therneau T. Mixed effects Cox models. CRAN repository. 2015 Jun 15;1030.

19. Balduzzi S, Rücker G, Schwarzer G. How to perform a meta-analysis with R: a practical tutorial. BMJ Mental Health. 2019 Nov;22(4):153–60.

20. Abd ElHafeez S, D’Arrigo G, Leonardis D, Fusaro M, Tripepi G, Roumeliotis S. Methods to analyze time to event data: the cox regression analysis. Oxidative medicine and cellular longevity. 2021;2021(1):1302811.

21. Higgins JP, Thompson SG. Quantifying heterogeneity in a meta analysis. Statistics in medicine. 2002 Jun 15;21(11):1539–58.

22. Sasson G, Kruger Ben-Shabat S, Seroussi E, Doron-Faigenboim A, Shterzer N, Yaacoby S, Berg Miller ME, White BA, Halperin E, Mizrahi I. Heritable bovine rumen bacteria are phylogenetically related and correlated with the cow’s capacity to harvest energy from its feed. MBio. 2017 Sep 6;8(4):10–128.

23. Luan S, Duersteler M, Galbraith EA, Cardoso FC. Effects of direct-fed Bacillus pumilus 8G-134 on feed intake, milk yield, milk composition, feed conversion, and health condition of pre-and postpartum Holstein cows. Journal of Dairy Science. 2015 Sep 1;98(9):6423–32.

24. AlZahal O, McGill H, Kleinberg A, Holliday JI, Hindrichsen IK, Duffield TF, McBride BW. Use of a direct-fed microbial product as a supplement during the transition period in dairy cattle. Journal of dairy science. 2014 Nov 1;97(11):7102–14.

25. Hiltz RL, Steelreath MR, Degenshein-Woods MN, Hung HC, Aguilar A, Nielsen H, Rezamand P, Laarman AH. Effects of Saccharomyces cerevisiae boulardii (CNCM I-1079) on feed intake, blood parameters, and production during early lactation. Journal of Dairy Science. 2023 Jan 1;106(1):187–201.

26. Nicola MS, Velasquez BE, Silva LR, Meireles M, Lopes L, Cappellozza BI, Londero US, Da Silva TC, Komninou ER, Del Pino FA, Rabassa VR. Effects of supplementation with Bacillus licheniformis 809 and Bacillus subtilis 810 on colostrum quality, productive performance, nutrient digestibility, and metabolic responses of dairy cows during the transition period. Journal of Dairy Science. 2026 Jan 20.

27. Overton TR, Waldron MR. Nutritional management of transition dairy cows: strategies to optimize metabolic health. Journal of dairy science. 2004 Jul 1;87:E105–19.

